# Role of Rare and Low Frequency Variants in Gene-Alcohol Interactions on Plasma Lipid Levels

**DOI:** 10.1101/561225

**Authors:** Zhe Wang, Han Chen, Traci M. Bartz, Lawrence F. Bielak, Daniel I. Chasman, Mary F. Feitosa, Nora Franceschini, Xiuqing Guo, Elise Lim, Raymond Noordam, Melissa A. Richard, Heming Wang, Brian Cade, L. Adrienne Cupples, Paul S. de Vries, Franco Giulanini, Jiwon Lee, Rozenn N. Lemaitre, Lisa W. Martin, Alex P. Reiner, Stephen S. Rich, Pamela J. Schreiner, Stephen Sidney, Colleen M. Sitlani, Jennifer A. Smith, Ko Willems van Dijk, Jie Yao, Wei Zhao, Myriam Fornage, Sharon L.R. Kardia, Charles Kooperberg, Ching-Ti Liu, Dennis O Mook-Kanamori, Michael A. Province, Bruce M. Psaty, Susan Redline, Paul M. Ridker, Jerome I. Rotter, Eric Boerwinkle, Alanna C. Morrison

**Affiliations:** Human Genetics Center, University of Texas Health Science Center at Houston, Houston, TX; Center for Precision Health, School of Public Health & School of Biomedical Informatics, University of Texas Health Science Center at Houston, Houston, TX; Cardiovascular Health Research Unit, Department of Biostatistics, University of Washington, Seattle, WA; School of Public Health, Department of Epidemiology, University of Michigan, Ann Arbor, MI; Division of Preventive Medicine, Brigham and Women’s Hospital, Boston, MA; Harvard Medical School, Boston, MA; Division of Statistical Genomics, Department of Genetics, Washington University School of Medicine, St. Louis, MO; Department of Epidemiology, Gillings School of Global Public Health, University of North Carolina, Chapel Hill, NC; The Institute for Translational Genomics and Population Sciences, Department of Pediatrics, Los Angeles Biomedical Research Institute at Harbor-UCLA Medical Center, Torrance, CA; Biostatistics Department, Boston University School of Public Health, Boston, MA; Section of Gerontology and Geriatrics, Department of Internal Medicine, Leiden University Medical Center, Leiden; Brown Foundation Institute of Molecular Medicine, The University of Texas Health Science Center at Houston, Houston, TX; Division of Sleep and Circadian Disorders, Department of Medicine, Brigham and Women’s Hospital, Boston, MA; NHLBI Framingham Heart Study, Framingham, MA; Cardiovascular Health Research Unit, Department of Medicine, University of Washington, Seattle, WA; George Washington University School of Medicine and Health Sciences, Washington, DC; Fred Hutchinson Cancer Research Center, Seattle, WA; Center for Public Health Genomics, Department of Public Health Sciences, University of Virginia, Charlottesville, VA; Epidemiology & Community Health, School of Public Health, University of Minnesota, Minneapolis, MN; Division of Research, Kaiser Permanente of Northern California, Oakland, CA; Institute for Social Research, Survey Research Center, University of Michigan, Ann Arbor, MI; Department of Human Genetics, Leiden University Medical Center, Leiden; Division of Endocrinology, Department of Internal Medicine, Leiden University Medical Center, Leiden; Department of Clinical Epidemiology, Leiden University Medical Center, Leiden; Department of Public Health and Primary Care, Leiden University Medical Center, Leiden; Cardiovascular Health Research Unit, Department of Epidemiology, University of Washington, Seattle, WA; Cardiovascular Health Research Unit, Department of Health Services, University of Washington, Seattle, WA; Kaiser Permanente Washington Health Research Institute, Seattle, WA; Human Genome Sequencing Center, Baylor College of Medicine, Houston, TX

**Keywords:** gene-environment interactions, lipid levels, alcohol consumption, genome-wide association study, rare variant test

## Abstract

**Background:** Alcohol intake influences plasma lipid levels and such effects may be modulated by genetic variants.

**Objective:** We aimed to characterize the role of aggregated rare and low-frequency variants in gene by alcohol consumption interactions associated with fasting plasma lipid levels.

**Design:** In the Cohorts for Heart and Aging Research in Genomic Epidemiology (CHARGE) consortium, fasting plasma triglycerides (TG), and high- and low-density lipoprotein cholesterol (HDL-c and LDL-c) were measured in 34,153 European Americans from five discovery studies and 32,275 individuals from six replication studies. Rare and low-frequency protein coding variants (minor allele frequency ≤ 5%) measured by an exome array were aggregated by genes and evaluated by a gene-environment interaction (GxE) test and a joint test of genetic main and GxE interaction effects. Two dichotomous self-reported alcohol consumption variables, current drinker, defined as any recurrent drinking behavior, and regular drinker, defined as the subset of current drinkers who consume at least two drinks per week, were considered.

**Results:** We discovered and replicated 21 gene-lipid associations at 13 known lipid loci through the joint test. Eight loci (*PCSK9, LPA, LPL, LIPG, ANGPTL4, APOB, APOC3 and CD300LG*) remained significant after conditioning on the common index single nucleotide polymorphism (SNP) identified by previous genome-wide association studies, suggesting an independent role for rare and low-frequency variants at these loci. One significant gene-alcohol interaction on TG was discovered at a Bonferroni corrected significance level (*p*-value <5×10^−5^) and replicated (*p*-value <0.013 for the interaction test) in *SMC5*.

**Conclusions:** In conclusion, this study applied new gene-based statistical approaches to uncover the role of rare and low-frequency variants in gene-alcohol consumption interactions on lipid levels.

## Introduction

Plasma lipid profiles, including high-density lipoprotein cholesterol (HDL-c), low-density lipoprotein cholesterol (LDL-c), and triglyceride (TG) levels have been well characterized for their roles in the development and prevention of cardiovascular disease (CVD) (1, 2). Genome-wide association studies (GWAS) and advanced DNA sequence technology have uncovered more than two hundred genetic loci influencing lipid levels (3–9), and these common (minor allele frequency [MAF] >5%) single nucleotide polymorphisms (SNPs) often reside in non-coding regions of the genome. In addition to the evidence that genetic factors affect plasma lipid profiles, environmental factors influence lipid levels as well. Epidemiologic studies have demonstrated an association between moderate alcohol consumption and improved lipid profile, including higher HDL-C levels, HDL particle concentration, and HDL-C subfractions (10, 11). However, the association between alcohol use and LDL-C or TG levels is unclear. Some studies reported positive associations while others reported negative associations (12–21).

Studying gene-by-environment (G×E) interactions is important, as it extends our knowledge of the genetic architecture of complex traits and improves our understanding of the underlying mechanisms of common diseases for novel and known loci (22–24). Several large-scale genome-wide G×E studies have successfully identified novel common variants accounting for the environmental effects such as alcohol consumption and smoking status on lipid levels and other CVD related traits (25–34). These studies have successfully identified common variant loci that were not detected via analysis of main effects alone. However, unlike well-established G×E interaction tests for common variants (35, 36), methods for detecting rare variant G×E interactions are emerging. Recently developed novel approaches for testing rare variant G×E interaction effects include a joint test that allows for simultaneous testing of the genetic main effect and interaction effect as well as the ability to assess gene-based GxE interactions (37).

Accounting for the effect of alcohol consumption in defining the genetic architecture of lipid levels may not only provide valuable insights into relationship between alcohol consumption and lipids, but also may help refine association signals at previously identified GWAS loci or identify new loci. This study is the first to incorporate G×E interaction in modeling rare and low-frequency variant genetic and alcohol effects on plasma lipid levels.

## Methods

### Overview of participating studies

This study includes 66,428 men and women between 18-80 years of age from 11 European-ancestry population studies that are part of the CHARGE Gene-Lifestyle Interactions Working Group (24). The participating studies include the Atherosclerosis Risk in Communities (ARIC) study, the Coronary Artery Risk Development in Young Adults (CARDIA) study, the Framingham Heart Study (FHS), the Netherlands Epidemiology of Obesity (NEO) study, the Women’s Health Initiative (WHI) study, the Cleveland Family Study (CFS), the Cardiovascular Health Study (CHS), the Family Heart Study (FamHS), the Genetic Epidemiology Network of Arteriopathy (GENOA) study, the Multi-Ethnic Study of Atherosclerosis (MESA), and the Women’s Genome Health Study (WGHS). Additional detail for these studies is provided in the Supplemental Materials. Each study obtained informed consent from participants and approval from the appropriate institutional review boards. A total of 34,153 participants from five studies participated in the discovery phase (ARIC, FHS, NEO, WHI and CARDIA), and six studies involving 32,275 participants were used for replication (WGHS, CFS, CHS, FamHS, GENOA and MESA).

### Plasma lipids and alcohol consumption

Three fasting (≥ 8 hours) lipid measures were analyzed separately. HDL-C and TG were directly assayed, while LDL-C was either directly assayed (WGHS, FamHS if TG > 400 mg/dL) or estimated using the Friedewald equation (38) (ARIC, FHS, NEO, WHI, CARDIA, CFS, CHS, FamHS, GENOA, MESA) in samples with TG ≤ 400 mg/dL. LDL-C levels were adjusted for use of statins: if LDL-C levels were directly assayed, LDL-C levels were adjusted for lipid-lowering medication use by dividing the original levels by 0.7, otherwise, LDL-C levels were adjusted by first dividing total cholesterol by 0.8, and then using the corrected total cholesterol level in the Friedewald equation. When information on statin-specific use was unavailable, LDL-C levels were adjusted for use of unspecified lipid-lowering medication, but only if lipid measurements were performed after 1994. Due to their skewed distributions, HDL-C and TG were natural log transformed prior to analyses.

Alcohol consumption was assessed using two dichotomized self-reported alcohol consumption variables: “current drinker” status, defined as any recurrent drinking behavior, and “regular drinker” status, as the subset of current drinkers who consume at least two drinks per week (33). For this study, definition of “a drink” is approximately 13g of pure ethanol, and this measure was used to standardize the definitions across studies.

### Genotyping and quality control

Genotyping was performed using the Illumina or Affymetrix Human Exome array v1 or v1.1. To improve accurate calling of rare variants, genotyped data from 10 CHARGE Consortium studies were jointly called (39). Using the curated clustering files from the CHARGE joint calling effort, several cohorts within our study re-called their genotypes. For the remainder of participating studies, genotypes were determined using either BeadStudio or Zcall (40). Detailed information regarding the genotyping platform for each study is presented in Supplemental Table 1. All studies performed the following sample-level quality control steps: call rate <95%, autosomal heterozygosity outliers, gender discordance, GWAS discordance (if GWAS data available), ethnic outlier in a principal components analysis. Variants were removed by filtering for Hardy-Weinberg equilibrium test *p*-value (pHWE) < 5×10^−6^, call rate <95%, and poorly clustering variants.

### Study-specific association analyses

Statistical analyses were performed within each study using the gene-based rareGE R package (37), and were performed for each lipid/alcohol consumption combination for a total of six combinations. Two types of analyses were considered: 1) a GxE test that considers the genetic main effects as fixed/random effects, and 2) a joint analysis of the genetic main and the GxE interaction effects. Rare and low-frequency (MAF ≤ 5%) functional variants (i.e. frameshift, nonsynonymous, stop/gain, stop/loss, and splicing) were aggregated within genes. Genes with 0 or only 1 rare and low-frequency variant, or genes with a cumulative minor allele count ≤ 10 were not analyzed within each study. Models were adjusted for age, sex, principal components (PCs) and additionally study-specific covariates as presented in Supplemental Table 1.

### Meta-analyses

A weighted Z-test using square root of sample sizes as weights was used to meta-analyze study-specific *p*-values for genes present in at least 2 discovery studies (41). Genes of interest from the discovery phase with a *p*-value < 5×10^−5^ were pursued for replication. For these select genes, we used the same approaches as in discovery studies to perform meta-analysis of the replication studies. Significance was determined using a Bonferroni correction for the number of gene-lipid pairs taken forward to replication (*p*-value < 0.05/30 = 0.0017 for analysis of the joint test of genetic main and interaction effects, *p*-value <0.05/4 = 0.013 for analysis of the interaction effects).

### Additional analyses: conditional and single variant tests

For each replicated gene-lipid pair, additional analyses were conducted following the flowchart shown in Figure 1. For genes +/-500kb bp from previously reported lipid loci (42), conditional analyses were performed to identify aggregated rare and low-frequency variants associated with lipids independent of the previously reported common index single nucleotide polymorphism (SNP) (42). Results from study-specific conditional analyses were meta-analyzed using a weighted Z-test, separately in discovery and replication. For novel genes and known genes that remained significant after conditional analyses, we performed single variant tests for each variant (MAF ≤ 5% and minor allele count ≥ 5) that was included in the aggregate test in order to identify the driving variants within these genes. We obtained robust estimates of covariance matrices and robust standard errors from each study and implemented METAL to jointly meta-analyze the genetic main and interaction effects (36, 43), and to meta-analyze the interaction coefficients alone using inverse-variance weighted meta-analysis for each single variant within selected genes.

**Figure 1.**
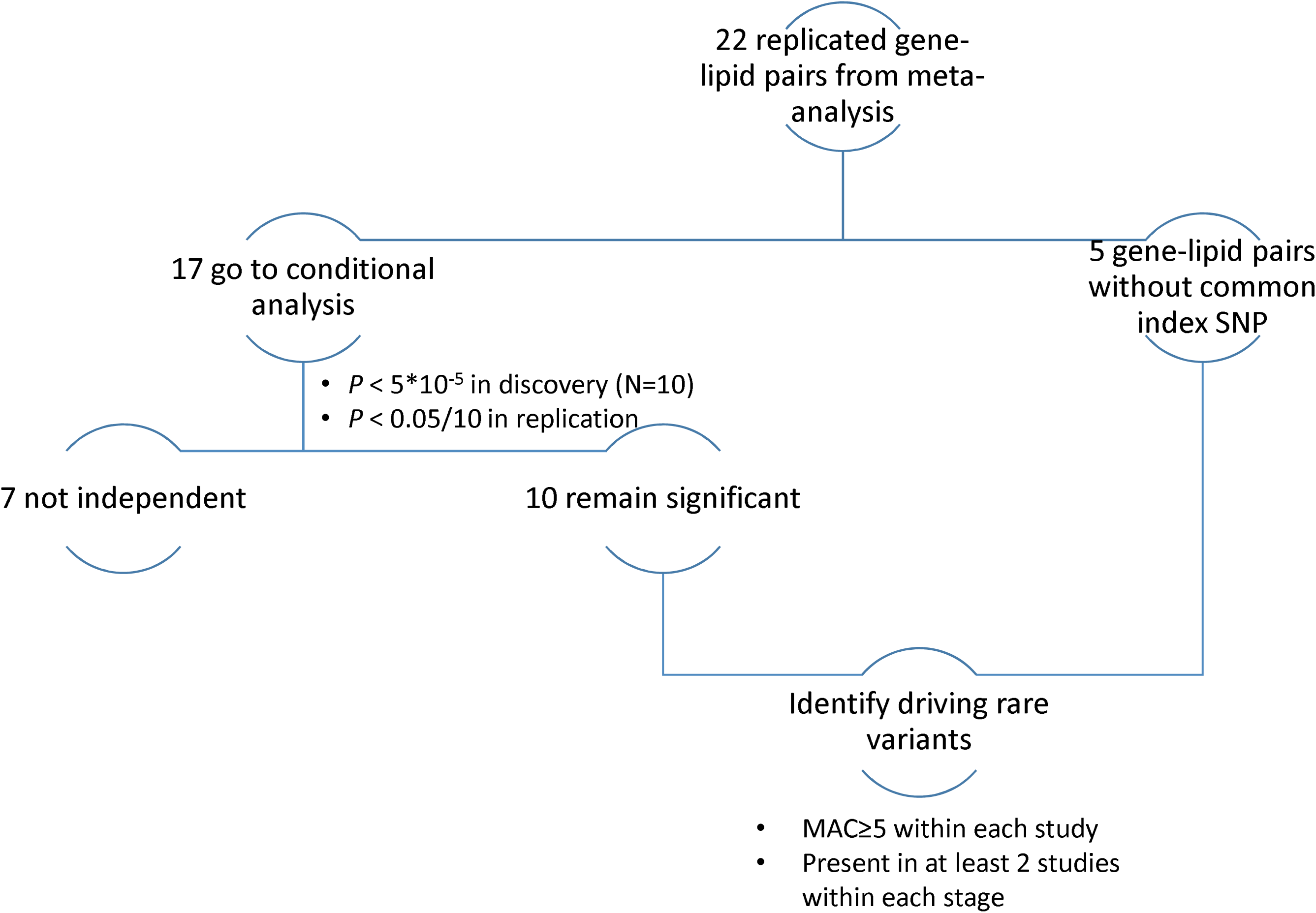
Flowchart of follow-up analyses, including conditional analysis and single variant test to identify driving rare variants For conditional analysis, significant results were defined as *p*-value < 5×10^−5^ in meta-analysis of discovery studies, and *p*-value < 0.05/10 (Bonferroni correction for 10 gene-lipid pairs with *p*- value < 5×10^−5^ in discovery phase) in meta-analysis of replication studies. For single variant test to identify driving rare variants, we applied Bonferroni correction for number of SNPs tested in discovery phase and number of SNPs taken forward to replication separately for joint test and interaction test for each lipid trait.

## Results

Descriptive statistics for the discovery and replication studies are summarized in Table 1 and Supplemental Table 1. On average, two thirds of the study participants were current drinkers and 39.5 percent were regular drinkers. The proportion of current and regular drinkers was greater for the discovery studies as compared to the replication studies.

**Table 1.** Descriptive characteristics for discovery and replication studies.

|  | **Study** | **Design** | **N** | **Curdrinker (%)** | **RegDrinker (%)** |
| --- | --- | --- | --- | --- | --- |
| Discovery | ARIC | Unrelated | 10989 | 64.9 | 36.8 |
|  | FHS | Family | 7258 | 83.6 | 65.5 |
|  | NEO | Unrelated | 5718 | 86.8 | 69.0 |
|  | WHI | Unrelated | 8021 | 76.5 | 32.6 |
|  | CARDIA | Unrelated | 2167 | 68.7 | 59.6 |
| Total/Average |  |  | 34,153 | 75.5 | 48.7 |
| Replication | WGHS | Unrelated | 22478 | 56.7 | 29.3 |
|  | CFS | Family | 253 | 50.2 | 25.1 |
|  | CHS | Unrelated | 3688 | 53.8 | 25.0 |
|  | FamHS | Family | 1735 | 50.7 | 28.3 |
|  | GENOA | Family | 1543 | 53.1 | 29.2 |
|  | MESA | Unrelated | 2578 | 71.9 | 43 |
| Total/Average |  |  | 32,275 | 57.2 | 29.8 |
| Overall |  |  | 66,428 | 66.6 | 39.5 |

Overall, meta-analysis showed highly consistent results across current drinker and regular drinker (Supplemental Table 2). Distributions of QQ plots for meta-analyzing discovery studies are shown in Supplemental Figure 1. In the discovery phase, we observed 31 gene-lipid associations (*p*-value < 5×10^−5^) in the joint analysis and 5 gene-lipid associations (*p*-value < 5×10^−5^) in the interaction test, with 3 genes overlapping between the two approaches (Supplemental Table 2). These gene-lipid pairs were taken forward for replication, one of which (*IDNK)* was only available in one replication study (the CHS). Therefore, we evaluated 30 gene-lipid associations for replication using the joint test and 4 using gene-alcohol interaction (Supplemental Table 2). Thirteen known lipid loci (21 gene-lipid associations) were replicated and one novel interaction were replicated for the *SMC5*-by-current drinker interaction on TG levels (Table 2). Among the replicated genes, 4 were shared between TG and HDL-C but none were shared between LDL-C and TG or HDL-C, as shown in a Venn diagram (Figure 2).

**Table 2.**
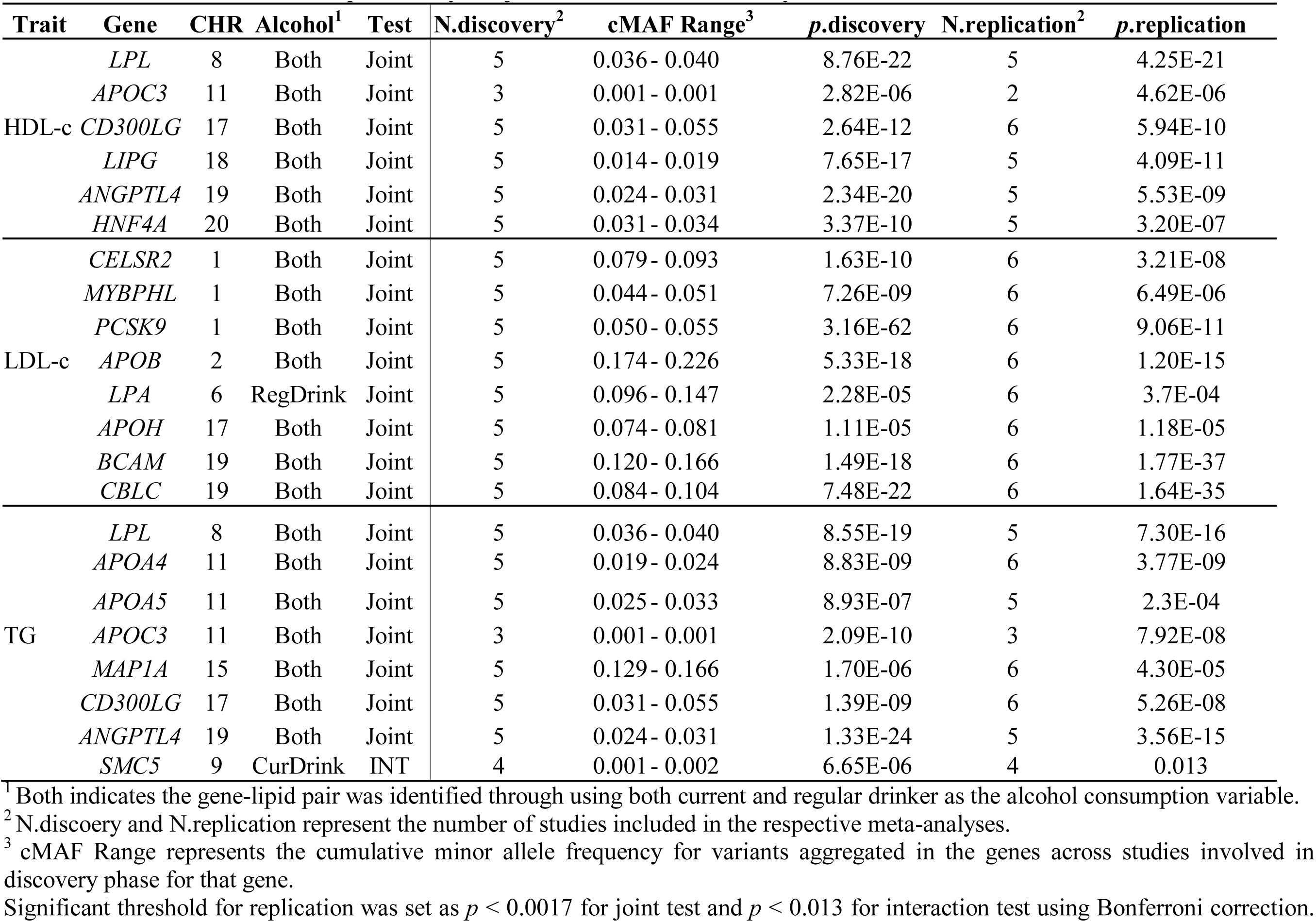
Genes discovered and replicated by the joint test or interaction only test.

**Figure 2.**
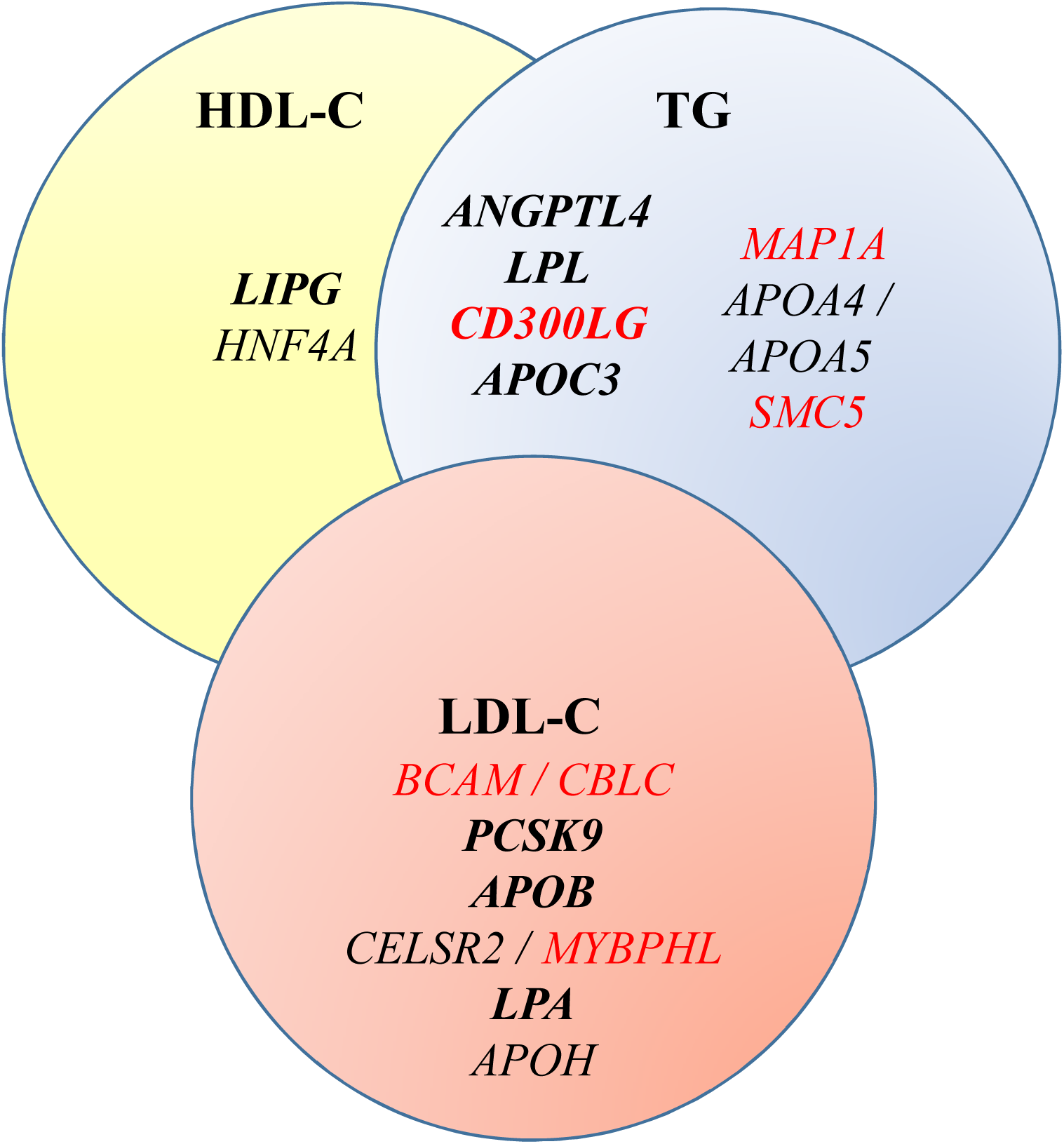
Genes as revealed by GxE interaction test or jointly testing the gene and GxE interaction effects in association with plasma lipid levels. **Bolded** genes were genes remained significant after conditioning on common index SNPs. Genes in red were not previously reported to be associated with one or more lipid traits

For the 13 known lipid loci that were replicated through the joint test, we performed conditional analyses in order to examine whether the gene-based rare variant effects are independent of the common index SNP identified by previous GWAS. In total, 8 loci (*PCSK9, LPA, LPL, LIPG, ANGPTL4, APOB, APOC3 and CD300LG*) (10 gene-lipid associations) remained significant after conditioning on a common index SNP. However, genes at known lipid loci yet these genes themselves were not previously reported to be associated with lipids, such as *BCAM* and *CBLC* on LDL-C, were strongly attenuated after adjusting for rs7412, the index SNPs of *APOE* identified by previous GWAS (Supplemental Table 3).

Single variant analyses were performed for the 5 gene-lipid associations that were not evaluated in the conditional analyses because they did not have previously reported common SNPs and for the 10 gene-lipid pairs that remained significant following conditional analyses (Figure 1, Supplemental Table 3). Single variant tests at these genes confirmed previous known low-frequency lipid variants. For example, rs11591147 in *PCSK9* was associated with LDL-C, and rs77960347 in *LIPG* and rs116843064 in *ANGPTL4* were associated with HDL-C. Additionally, we provide evidence that two of the driving variants underlying the joint test results are novel rare variants associated with LDL-c (Supplemental Table 4). One of them is rs41267813, a variant in the *LPA* gene (*p* = 6.55×10^−29^ discovery, *p* = 1.83×10^−03^ replication) and the other is rs41288783 of *APOB* gene (*p* = 5.40×10^−08^ discovery, *p* = 7.92×10^−07^ replication). For the novel interaction between *SMC5* and current drinker on TG levels, we identified the driving variant as rs142488686, a missense mutation (MAC = 5-7 discovery (ARIC and CARDIA), MAC = 7-17 replication (WGHS, CHS and MESA)), with positive interaction effect (*p* = 0.016 discovery, *p* = 0.008 replication), while the genetic main effect was modest (*p* < 0.1 discovery and replication, respectively).

## Discussion

This is the first large-scale study to evaluate the role of rare and low frequency variants in lipids by incorporating gene-alcohol consumption interactions. We tested for gene-alcohol interaction effect on lipid levels as well as the joint effects of genetic main and gene-alcohol interactions. We replicated 13 gene-lipid associations at known lipid loci, among which 2 leading rare variants in *APOB* and *LPA* genes associated with LDL-c were novel. Only one novel gene-alcohol interaction was identified as significant and successfully replicated (the interaction between rare and low-frequency variants in *SMC5* and current drinker on TG levels).

Using a single variant test, we confirmed previously identified rare and low-frequency lipid variants. For example, rs11591147 of *PCSK9* has been associated with LDL-c levels (44), rs77960347 of *LIPG* and rs116843064 of *ANGPTL4* have been associated with HDL-c levels (45, 46). A loss of function mutation in the *APOC3* gene, rs147210663, has been associated with a more than 40% lower average triglyceride level in individuals carrying one A allele (47, 48). In the present study, we observed a novel relationship between increased HDL-c levels in individuals carrying rs147210663 (A) allele as rs147210663 was previously reported as a founder mutation in a Pennsylvania Amish population (49).

Between the two novel rare driving variants we identified and replicated, rs41267813 (*LPA)* is located close to a stop/gain variant rs41267811 *(LPA)* that was also significantly associated with LDL-c levels in the discovery phase. However, we were unable to replicate the association with rs41267811 as it was only available in one replication study (WGHS) and therefore did not meet our criteria to be included in replication. *LPA* encoded protein constitutes a substantial portion of lipoprotein(a) and associated with inherited conditions including type III hyperlipoproteinemia and familial hyperlipidemia (50). A stop/gain mutation in this gene would be associated with lower LDL-C levels in carriers, which is true among non-drinkers. However, such effect may be modified by alcohol consumption as we observed the carriers of this variant with a higher LDL-C levels compared to non-carriers in a population who had at least two drinks per week in the ARIC study. Previous studies have reported a relationship between moderate alcohol consumption and lower Lp(a) lipoprotein concentrations (51, 52), but there was no existing evidence linking genetic variants of *LPA* to alcohol consumption. Although the underlying biology of the observed modification effect of alcohol consumption on rs41267813 and LDL-c associations remained unclear, we hypothesize that alcohol modifies the *LPA* expression for carriers of rs41267813, therefore modifies LDL-c levels.

In addition to the variant described above, the other driving rare variant had not been previously associated with a lipid trait, rs41288783 (p.Pro994Leu), a deleterious variant in *APOB* gene. A previous study reported its existence in a patient who was clinically diagnosed as familial hypercholesterolaemia (FH) without a detectable mutation (53). FH is characterized by very high levels of LDL-c, and we observed an association with higher LDL-c levels though jointly testing the effects of rs41288783 and its interaction with alcohol consumption. Nevertheless, the exact biological function of rs41288783 remains unknown. We note that a Mendelian randomization study has suggested a causal role of alcohol consumption in reducing plasma apo B and LDL-c levels in a general population (54). Considering this, alcohol consumption may have contributed to the observed significant joint effect of *APOB* and alcohol consumption on LDL-c levels.

For the significant gene-alcohol interaction effect we observed on TG levels, the driving variant was identified as rs142488686, a missense mutation in *SMC5* (Structural Maintenance Of Chromosomes 5). *SMC5* encodes a core component involved in repair of DNA double-strand breaks and required for telomere maintenance (55–57). Variants in *SMC5* have not been previously reported to be associated with lipid levels nor alcohol consumption, and it is unknown whether the interaction between *SMC5* locus and current drinking behavior on TG levels has a biological aspect.

A limitation of this study is the imbalance in percentage of alcohol consumers between discovery (on average 48.7% regular drinker, 78.5% current drinker) and replication studies (on average 29.8% regular drinker, 57.2% current drinker) which may have impacted our ability to identify and replicate additional loci beyond what is reported here. Additionally, as self-reported alcohol consumption was used and may very likely be underreported, this study may suffer from loss of statistical power due to potential misclassification (58). Similarly, dichotomizing alcohol consumption into regular drinkers and current drinkers may also reduce power as compared to treating it as a continuous variable (59). It is possible that a more comprehensive characterization of alcohol consumption could reveal associations that were missed in the present study. In addition, although the sample size of 66,428 may seem sufficient for a traditional GWAS, to identify additional novel loci while focusing on rare variants and gene-environment interactions may require larger sample size or bigger effect size (23, 60).

In conclusion, this study applied emerging statistical approaches to investigate the role of rare and low-frequency variants in gene-alcohol consumption interaction effects on lipid levels, and identified 2 novel rare variants at know lipid loci for LDL-c levels and 1 novel gene-alcohol interaction for TG levels. Our results show promise for other larger scale studies analyzing rare variant GxE interactions to refine association signals at previously identified loci to reveal novel biology.

## Supporting information

Supplemental Materials_acknowledgement

## Acknowledgments

The various Gene-Lifestyle Interaction projects, including this one, are largely supported by a grant from the U.S. National Heart, Lung, and Blood Institute (NHLBI), the National Institutes of Health, R01HL118305. Full set of study-specific funding sources and acknowledgments appear in the Supplemental Materials.

## Authors’ Contributions

The authors’ contributions were as follows – ZW, DIC, RN, HW, LAC, KWvD., JAS, SSR, MF, SLRK, CL, DOM, MAP, PMR, JIR, EB, and ACM: designed research (project conception, development of overall research plan, and study oversight); LFB, DIC, MFF, NF, XG, BC, KWvD, JL, LWM, APR, SSR, PJS, SS, JAS, WZ, SLRK, CK, DOM, MAP, BMP, SR, PMR, and JIR: conducted research (hands-on conduct of the experiments and data collection) and provided essential reagents or provided essential materials (applies to authors who contributed by providing animals, constructs, databases, etc, necessary for the research); WZ, HC, TMB, LFB, DIC, MFF, NF, XG, EL, RN, MAR, HW, LAC, FG, KWvD, JL, JY, MF, SLRK, CL, DOM, BMP, and JIR: analyzed data or performed statistical analysis; ZW and ACM: wrote the paper; and all authors: read, reviewed, and approved of the final manuscript.

