## Supplemental Materials_acknowledgement for "Role of Rare and Low Frequency Variants in Gene-Alcohol Interactions on Plasma Lipid Levels"

### **Discovery Study Description:**

**ARIC (Atherosclerosis Risk in Communities):** The ARIC study is a population-based prospective cohort study of cardiovascular disease sponsored by the National Heart, Lung, and Blood Institute (NHLBI). ARIC included 15,792 individuals, predominantly European American and African American, aged 45-64 years at baseline (1987-89), chosen by probability sampling from four US communities. Cohort members completed three additional triennial follow-up examinations and a fifth exam in 2011-2013. The ARIC study has been described in detail previously (1). We only analyzed the European Americans for this project.

**FHS (Framingham Heart Study):** FHS began in 1948 with the recruitment of an original cohort of 5,209 men and women (mean age 44 years; 55 percent women). In 1971 a second generation of study participants was enrolled; this cohort (mean age 37 years; 52% women) consisted of 5,124 children and spouses of children of the original cohort. A third generation cohort of 4,095 children of offspring cohort participants (mean age 40 years; 53 percent women) was enrolled in 2002-2005 and are seen every 4 to 8 years. Details of study designs for the three cohorts are summarized elsewhere. At each clinic visit, a medical history was obtained with a focus on cardiovascular content, and participants underwent a physical examination including measurement of height and weight from which BMI was calculated.

**NEO (The Netherlands Epidemiology of Obesity study):** The NEO was designed for extensive phenotyping to investigate pathways that lead to obesity-related diseases. The NEO study is a population-based, prospective cohort study that includes 6,671 individuals aged 45–65 years, with an oversampling of individuals with overweight or obesity. At baseline, information on demography, lifestyle, and medical history have been collected by questionnaires. In addition, samples of 24-h urine, fasting and postprandial blood plasma and serum, and DNA were collected. Genotyping was performed using the Illumina HumanCoreExome chip, which was subsequently imputed to the 1000 genome reference panel. Participants underwent an extensive physical examination, including anthropometry, electrocardiography, spirometry, and measurement of the carotid artery intima-media thickness by ultrasonography. In random subsamples of participants, magnetic resonance imaging of abdominal fat, pulse wave velocity of the aorta, heart, and brain, magnetic resonance spectroscopy of the liver, indirect calorimetry, dual energy X-ray absorptiometry, or accelerometry measurements were performed. The collection of data started in September 2008 and completed at the end of September 2012. Participants are currently being followed for the incidence of obesity-related diseases and mortality.

**WHI (Women’s Health Initiative):** WHI is a long-term national health study that focuses on strategies for preventing common diseases such as heart disease, cancer and fracture in postmenopausal women. A total of 161,838 women aged 50–79 years old were recruited from 40 clinical centers in the US between 1993 and 1998. WHI consists of an observational study, two clinical trials of postmenopausal hormone therapy (HT, estrogen alone or estrogen plus progestin), a calcium and vitamin D supplement trial, and a dietary modification trial.(2) Study recruitment and exclusion criteria have been described previously. Recruitment was done through mass mailing to age-eligible women obtained from voter registration, driver’s license and Health Care Financing Administration or other insurance list, with emphasis on recruitment of minorities and older women.(3) Exclusions included participation in other randomized trials, predicted survival < 3 years, alcoholism, drug dependency, mental illness and dementia. For the CT, women were ineligible if they had a systolic BP > 200 mm Hg or diastolic BP > 105 mm Hg, a history of hypertriglyceridemia or breast cancer. Study protocols and consent forms were approved by the IRB at all participating institutions. Socio-demographic characteristics, lifestyle, medical history and self-reported medications were collected using standardized questionnaires at the screening visit. Physical measures of height, weight and blood pressure were measured at a baseline clinical visit.(3) BP was measured by certified staff using standardized procedures and instruments.(4) Two BP measures were recorded after 5 minutes rest using a mercury sphygmomanometer. Appropriate cuff bladder size was determined at each visit based on arm circumference. Diastolic BP was taken from the phase V Korotkoff measures. The average of the two measurements, obtained 30 seconds apart, was used in analyses.

**CARDIA (Coronary Artery Risk Development in Young Adults):** CARDIA is a prospective multicenter study with 5,115 adults Caucasian and African American participants of the age group 18-30 years, recruited from four centers at the baseline examination in 1985-1986. The recruitment was done from the total community in Birmingham, AL, from selected census tracts in Chicago, IL and Minneapolis, MN; and from the Kaiser Permanente health plan membership in Oakland, CA. The details of the study design for the CARDIA study have been previously published. Nine examinations have been completed since initiation of the study, respectively in the years 0, 2, 5, 7, 10, 15, 20, 25, and 30. Written informed consent was obtained from participants at each examination and all study protocols were approved by the institutional review boards of the participating institutions. We only analyzed the European Americans for this project.

### **Replication Study Description**

**WGHS (Women’s Genome Health Study):** WGHS is a prospective cohort of female North American health care professionals representing participants in the Women’s Health Study (WHS) trial who provided a blood sample at baseline and consent for blood-based analyses. Participants in the WHS were 45 years or older at enrollment and free of cardiovascular disease, cancer or other major chronic illness. The current data are derived from 23,294 WGHS participants for whom whole genome genotype information was available at the time of analysis and for whom self-reported European ancestry could be confirmed by multidimensional scaling analysis of 1,443 ancestry informative markers in PLINK v. 1.06. At baseline, BP and lifestyle habits related to smoking, consumption of alcohol, and physical activity as well as other general clinical information were ascertained by a self-reported questionnaire, an approach which has been validated in the WGHS demographic, namely female health care professionals.

**CFS (Cleveland Family Study):** The Cleveland Family Study is a family-based, longitudinal cohort study with the objective of examining the genetic and familial basis of sleep apnea. It consists of 2534 African- and European-American individuals from 356 families. Index probands with confirmed sleep apnea were recruited from northern Ohio sleep centers. Up to four visits were made from 1990 to 2006, and a final laboratory visit at a clinical research center between 2000 and 2006. Blood samples were obtained from 1447 participants from the final two exams, and DNA was extracted from samples that passed quality control. In this analysis, we subsetted to only European Americans who had a lab visit, and were 18 or older at the time of visit.

**CHS (Cardiovascular Health Study):** CHS is a population-based cohort study of risk factors for cardiovascular disease in adults 65 years of age or older conducted across four field centers.(5) The original predominantly European ancestry cohort of 5,201 persons was recruited in 1989-1990 from random samples of the Medicare eligibility lists and an additional predominately African-American cohort of 687 persons was enrolled in 1992-93 for a total sample of 5,888. Blood samples were drawn from all participants at their baseline examination and DNA was subsequently extracted from available samples. CHS was approved by institutional review committees at each site and individuals in the present analysis gave informed consent including consent to use of genetic information for the study of cardiovascular disease. We only analyzed the European Americans for this project.

**FamHS (Family Heart Study):** The NHLBI FamHS study design, collection of phenotypes and covariates as well as clinical examination have been previously described (<https://dsgweb.wustl.edu/fhscc/>) (6). In brief, the FamHS recruited 1,200 families (approximately 6,000 individuals), half randomly sampled, and half selected because of an excess of coronary heart disease (CHD) or risk factor abnormalities as compared with age- and sex-specific population rates. The participants were sampled from four population-based parent studies: the Framingham Heart Study, the Utah Family Tree Study, and two centers for the Atherosclerosis Risk in Communities study (ARIC: Minneapolis, and Forsyth County, NC). These individuals attended a clinic exam (1994-1996) and a broad range of phenotypes were assessed in the general domains of CHD, atherosclerosis, cardiac and vascular function, inflammation and hemostasis, lipids and lipoproteins, blood pressure, diabetes and insulin resistance, pulmonary function, diet, education, socioeconomic status, habitual behavior, physical activity, anthropometry, medical history and medication use. Approximately 8 years later, study participants belonging to the largest pedigrees were invited for a second clinical exam (2002-04). The most important CHD risk factors were measured again, including lipids, parameters of glucose metabolism, blood pressure, anthropometry, and several biochemical and hematologic markers. In addition, a computed tomography examination provided measures of coronary and aortic calcification, and abdominal and liver fat burden. Medical history and medication use was updated. A total of 2,756 European ancestry subjects in 510 extended random and high CHD risk families were studied.

**GENOA (Genetic Epidemiology Network of Arteriopathy):** GENOA is one of four networks in the NHLBI Family-Blood Pressure Program (FBPP) (7, 8). GENOA's long-term objective is to elucidate the genetics of target organ complications of hypertension, including both atherosclerotic and arteriolosclerotic complications involving the heart, brain, kidneys, and peripheral arteries. The longitudinal GENOA Study recruited European-American and African-American sibships with at least 2 individuals with clinically diagnosed essential hypertension before age 60 years. All other members of the sibship were invited to participate regardless of their hypertension status. Participants were diagnosed with hypertension if they had either 1) a previous clinical diagnosis of hypertension by a physician with current anti-hypertensive treatment, or 2) an average systolic blood pressure ≥ 140 mm Hg or diastolic blood pressure ≥ 90 mm Hg based on the second and third readings at the time of their clinic visit. Exclusion criteria were secondary hypertension, alcoholism or drug abuse, pregnancy, insulin-dependent diabetes mellitus, or active malignancy. During the first exam (1995-2000), 1,583 European Americans from Rochester, MN and 1,854 African Americans from Jackson, MS were examined. Between 2000 and 2005, 1,241 of the European Americans and 1,482 of the African Americans returned for a second examination. We only analyzed the European Americans for this project.

**MESA (Multi-Ethnic Study of Atherosclerosis):** The Multi-Ethnic Study of Atherosclerosis (MESA) is a study of the characteristics of subclinical cardiovascular disease and the risk factors that predict progression to clinically overt cardiovascular disease or progression of the subclinical disease. MESA consisted of a diverse, population-based sample of an initial 6,814 asymptomatic men and women aged 45-84. 38 percent of the recruited participants were white, 28 percent African American, 22 percent Hispanic, and 12 percent Asian, predominantly of Chinese descent.(9) Participants were recruited from six field centers across the United States: Wake Forest University, Columbia University, Johns Hopkins University, University of Minnesota, Northwestern University and University of California - Los Angeles. Participants are being followed for identification and characterization of cardiovascular disease events, including acute myocardial infarction and other forms of coronary heart disease (CHD), stroke, and congestive heart failure; for cardiovascular disease interventions; and for mortality. The first examination took place over two years, from July 2000 - July 2002. It was followed by four examination periods that were 17-20 months in length. Participants have been contacted every 9 to 12 months throughout the study to assess clinical morbidity and mortality. We only analyzed the European Americans for this project.

### **Discovery Study Acknowledgement**

**ARIC (Atherosclerosis Risk in Communities):** The Atherosclerosis Risk in Communities study has been funded in whole or in part with Federal funds from the National Heart, Lung, and Blood Institute, National Institutes of Health, Department of Health and Human Services (contract numbers HHSN268201700001I, HHSN268201700002I, HHSN268201700003I, HHSN268201700004I and HHSN268201700005I). The authors thank the staff and participants of the ARIC study for their important contributions. Funding support for “Building on GWAS for NHLBI-diseases: the U.S. CHARGE consortium” was provided by the NIH through the American Recovery and Reinvestment Act of 2009 (ARRA) (5RC2HL102419).

**FHS (Framingham Heart Study):** This research was conducted in part using data and resources from the Framingham Heart Study of the National Heart Lung and Blood Institute of the National Institutes of Health and Boston University School of Medicine. The analyses reflect intellectual input and resource development from the Framingham Heart Study investigators participating in the SNP Health Association Resource (SHARe) project. This work was partially supported by the National Heart, Lung and Blood Institute's Framingham Heart Study (Contract Nos. N01-HC-25195 and HHSN268201500001I) and its contract with Affymetrix, Inc for genotyping services (Contract No. N02-HL-6-4278). A portion of this research utilized the Linux Cluster for Genetic Analysis (LinGA-II) funded by the Robert Dawson Evans Endowment of the Department of Medicine at Boston University School of Medicine and Boston Medical Center. This research was partially supported by grant R01-DK089256 from the National Institute of Diabetes and Digestive and Kidney Diseases (MPIs: Michael Province, L. Ching-Ti Liu, Kari North) and training grant T32GM074905-14 from NIH/National Institute of General Medical Sciences.

**NEO (The Netherlands Epidemiology of Obesity study):** The authors of the NEO study thank all individuals who participated in the Netherlands Epidemiology in Obesity study, all participating general practitioners for inviting eligible participants and all research nurses for collection of the data. We thank the NEO study group, Pat van Beelen, Petra Noordijk and Ingeborg de Jonge for the coordination , lab and data management of the NEO study. The genotyping in the NEO study was upported by the Centre National de Génotypage (Paris, France), headed by Jean Francois Deleuze. The NEO study is supported by the participating Departments, the Division and the Board of Directors of the Leiden University Medical Center, and by the Leiden University, Research Profile Area Vascular and Regenerative Medicine. Dennis Mook Kanamori is supported by Dutch Science Organization (ZonMW) VENI Grant 916.14.023). Diana van Heemst and Raymond Noordam were supported by the European Commission funded project HUMAN (Health-2013-INNOVATION-1-602757)

**WHI (Women’s Health Initiative):** The WHI program is funded by the National Heart, Lung, and Blood Institute through contracts HHSN268201600018C, HHSN268201600001C, HHSN268201600002C, HHSN268201600003C, and HHSN268201600004C. We thank the WHI investigators and staff for their dedication, and the study participants for making the program possible. A full listing of WHI investigators can be found at:

http://www.whi.org/researchers/Documents%20%20Write%20a%20Paper/WHI%20Investig

ator%20Long%20List.pdf

**CARDIA (Coronary Artery Risk Development in Young Adults):** The CARDIA Study is conducted and supported by the National Heart, Lung, and Blood Institute in collaboration with the University of Alabama at Birmingham (HHSN268201300025C & HHSN268201300026C), Northwestern University (HHSN268201300027C), University of Minnesota (HHSN268201300028C), Kaiser Foundation Research Institute (HHSN268201300029C), and Johns Hopkins University School of Medicine (HHSN268200900041C). CARDIA is also partially supported by the Intramural Research Program of the National Institute on Aging. Exome Chip genotyping was supported from grants R01-HL093029 and U01- HG004729 to MF.

### **Replication Study Acknowledgement**

**WGHS (Women’s Genome Health Study)**: The WGHS is supported by the National Heart, Lung, and Blood Institute (HL043851 and HL080467) and the National Cancer Institute (CA047988 and UM1CA182913), with funding for lipid fractions provided by the Donald W. Reynolds Foundation, and funding for genotyping provided by Amgen.

**CFS (Cleveland Family Study):** The CFS was supported by NHLBI R35HL135818, R01HL113338, R01HL098433, R01HL46380.

**CHS (Cardiovascular Health Study):** This CHS research was supported by NHLBI contracts HHSN268201200036C, HHSN268200800007C, HHSN268201800001C, N01HC55222, N01HC85079, N01HC85080, N01HC85081, N01HC85082, N01HC85083, N01HC85086; and NHLBI grants R01HL068986, U01HL080295, R01HL087652, R01HL105756, R01HL103612, R01HL120393, and U01HL130114 with additional contribution from the National Institute of Neurological Disorders and Stroke (NINDS). Additional support was provided through R01AG023629 from the National Institute on Aging (NIA). A full list of principal CHS investigators and institutions can be found at [CHS-NHLBI.org](http://chs-nhlbi.org/). The provision of genotyping data was supported in part by the National Center for Advancing Translational Sciences, CTSI grant UL1TR001881, and the National Institute of Diabetes and Digestive and Kidney Disease Diabetes Research Center (DRC) grant DK063491 to the Southern California Diabetes Endocrinology Research Center. The content is solely the responsibility of the authors and does not necessarily represent the official views of the National Institutes of Health.

**FamHS (Family Heart Study):** The FamHS was funded by R01HL118305 and R01HL117078 NHLBI grants, and 5R01DK07568102 and 5R01DK089256 NIDDK grant.

**GENOA (Genetic Epidemiology Network of Arteriopathy):** Support for GENOA was provided by the National Heart, Lung and Blood Institute (HL119443, HL118305, HL054464, HL054457, HL054481, HL071917 and HL087660) of the National Institutes of Health. Genotyping was performed at the Mayo Clinic (Stephen T. Turner, MD, Mariza de Andrade PhD, Julie Cunningham, PhD). We thank Eric Boerwinkle, PhD and Megan L. Grove from the Human Genetics Center and Institute of Molecular Medicine and Division of Epidemiology, University of Texas Health Science Center, Houston, Texas, USA for their help with genotyping. We would also like to thank the families that participated in the GENOA study.

**MESA (Multi-Ethnic Study of Atherosclerosis):** MESA and the MESA SHARe project are conducted and supported by the National Heart, Lung, and Blood Institute (NHLBI) in collaboration with MESA investigators. Support for MESA is provided by contracts HHSN268201500003I, N01-HC-95159, N01-HC-95160, N01-HC-95161, N01-HC-95162, N01-HC-95163, N01-HC-95164, N01-HC-95165, N01-HC-95166, N01-HC-95167, N01-HC-95168, N01-HC-95169, UL1-TR-000040, UL1-TR-001079, UL1-TR-001420, N02-HL-64278, UL1-TR-001881, and DK063491.
